# A Metabolomic Study of Cervical Dystonia

**DOI:** 10.1101/2020.08.30.274126

**Authors:** Chang Liu, Laura Scorr, Gamze Kilic-Berkmen, Adam Cotton, Stewart A. Factor, Alan Freeman, ViLinh Tran, Ken Liu, Karan Uppal, Dean Jones, H. A. Jinnah, Yan V. Sun

## Abstract

**Background:** Cervical dystonia is the most common of the adult-onset focal dystonias. Most cases are idiopathic. The current view is that cervical dystonia may be caused by some combination of genetic and environmental factors. Genetic contributions have been studied extensively, but there are few studies of other factors.

**Objective:** To conduct an exploratory metabolomics analysis of cervical dystonia to identify potentially abnormal metabolites or altered biological pathways.

**Methods:** Plasma samples from 100 cases with idiopathic cervical dystonia and 100 controls were compared using liquid chromatography coupled with mass spectrometry-based metabolomics.

**Results:** A total of 7,346 metabolic features remained after quality control, and 289 demonstrated significant differences between cases and controls, depending on statistical criteria chosen. Pathway analysis revealed 9 biological processes to be significantly associated at p<0.05, 5 pathways were related to carbohydrate metabolism, 3 pathways were related to lipid metabolism.

**Conclusion:** This is the first large scale metabolomics study for any type of dystonia. The results may provide potential novel insights into the biology of cervical dystonia.

## Introduction

Cervical dystonia (CD) is characterized by excessive involuntary contractions of muscles of the neck and shoulder leading to abnormal head postures, involuntary movements including tremor, and pain [1]. It is considered to be a rare disorder with an overall incidence of 1.2 per 100,000 person years [2], and crude prevalence of 29.5 per 100,000 population [3]. It typically begins in adults between 40-60 years of age, with females being affected more often than males [4]. For the majority of cases, a cause cannot be found, even after extensive diagnostic testing. Much of the recent research regarding potential causes has focused on genetics, as ∼15% of patients with CD have a family member affected with dystonia [4]. However, the genes currently known to cause dystonia account for no more than 4% of all cases [1]. These observations suggest that other, as yet undiscovered, genes may play a role.

Non-genetic factors may also have a place in pathogenesis. Rare cases of CD may be caused by focal lesions of the nervous system [5], exposure to medications such as neuroleptics [6, 7], infection [8], or autoimmune mechanisms [9]. A few small studies have methodically addressed other potential environmental contributions. A survey study of 184 cases of all types of idiopathic dystonia in Australia suggested cigarette smoking to be a risk factor [10]. A case control study of 67 cases of CD suggested a role for trauma to the head and neck [11].

The metabolome is the global collection of all small molecules in the body. It reflects the combined systemic effects of genetics, lifestyle, environmental exposure and secondary biological responses. Metabolomics, or the study of global metabolism, is an emerging discipline that has the potential to transform the study of the mechanism of disease development. High-resolution chemical profiling involves the use of liquid chromatography to separate analytes and high-resolution mass spectrometry to accurately measure mass and abundance [12, 13]. To our knowledge, there are no published metabolomic studies for any of the common adult-onset focal dystonia. The current study describes results of a blood-based metabolomics study of 100 subjects with CD and 100 healthy controls.

## Methods

### Study Population

The study was approved by the Emory Institutional Review Board, and all cases and controls provided written informed consent to participate. Subjects with a diagnosis of idiopathic CD were recruited from the Emory University Movement disorders clinic. Dystonia in other body regions was allowed (segmental and multifocal patterns), but involvement of neck muscles had to be the primary problem. Subjects were excluded if they had CD secondary to a known cause such as focal lesions, Parkinsonism or neuroleptic exposure. Control subjects were recruited from the same clinic. The majority of the controls were spouses or family members of the subjects with CD. Precise matching of cases and controls was not feasible.

We also collected information known to influence the metabolome. Subjects were asked if they use nicotine products currently or in the past or if they had a history of metabolic diseases (e.g., hypertension, hyperlipidemia, heart disease, or diabetes). Additionally, data about subjects’ body mass index (BMI) and diet were also collected.

### Untargeted High-Resolution Metabolic Profiling

We conducted high-resolution metabolic profiling using liquid chromatography with ultra-high resolution mass spectrometry (LC-HRMS; Thermo Scientific Fusion) following established protocols[14] and published methods[15, 16]. Plasma specimens were collected and stored at −80°C before thawing for liquid-chromatography mass spectrometry (LC/MS). Sam were treated with acetonitrile containing internal isotopic standard mix, centrifuged for 10 minutes at 4°C to remove protein, and then maintained in a Dionex Ultimate 3000 autosampler at 4°C until analysis. NIST 1950 [17] was analyzed at the beginning and end of the entire analysis procedure. Two replicated pooled human plasma samples were analyzed at the beginning, middle, and end of each batch of 40 study samples for batch effect correction and normalization. Samples were analyzed in triplicate using C18 chromatography (Higgins endcapped C18 2.1 cm × 5 cm × 3 µm column) the following gradient program: initial 0.5 min period of 60% A (LCMS grade water), 35% B (LCMS grade acetonitrile, 5% C (10 mM ammonium acetate), followed by linear increase to 0% A, 95% B, 5% C at 1.5 min which was held for an additional 3 min. Finally, mobile phase flow rate was held at 0.4 mL/min for 1.5 min, and then increased to 0.5 mL/min. MS was operated in negative ionization mode using electrospray ionization, mass resolution of 120,000 and data collected from 85 to 1275 m/z over 5 minutes)[15]. Source tune settings included capillary temperature, sheath gas, auxiliary gas, sweep gas and spray voltage settings of 300 °C, 45 (arbitrary units), 25 (arbitrary units), 1 (arbitrary units) and + 3.5 kV for negative mode. S-Lens RF level was maintained at 45. Feature extraction and alignment was performed in apLCMS [18], with quality control and feature filtering using xMSanalyzer [19]. Features are defined by their accurate mass *m/z*, retention time and ion abundance. Features with median coefficient of variation within technical replicates ≥ 75%, and samples with Pearson correlation coefficient ≥ 0.7 within triplicates were retained for downstream analysis. Batch effect was corrected using ComBat [20].

### Feature Annotation and Identification

Feature annotation was performed using xMSannotator [21] with the Human Metabolome Database [22]. Annotations with confidence score of 3 indicates high confidence level, score of 2 indicates medium confidence level. Only annotations with medium confidence or above were used. Furthermore, metabolite identification and confirmation was performed using an in-house library of metabolites that have been previously confirmed using comparison of retention time and MS/MS with authentic standards and comparison of MS/MS fragments with online spectral databases. The library was matched to the metabolic features that significantly associated with CD, allowing an m/z difference of 5 ppm and a retention time difference of 20 seconds. Feature identification levels were assigned according to the published criteria [23]. At level 1, the proposed structure was confirmed via appropriate measurement of a reference standard with MS, MS/MS and retention time matching. At level 2, the proposed structure was confirmed by MS/MS and matched with online databases or in-silico predicted spectra. At level 3, the proposed structure was confirmed by MS/MS at the chemical class level, but no evidence for a specific metabolite. At level 4, the proposed structure was computationally annotated using xMSannotator with confidence level medium or above [21]. At level 5, the proposed structure was matched accurately by mass.

### Statistical Analysis

Subject characteristics were reported as mean (standard deviation) for continuous variables and frequency (percentage) for categorical variables. The differences between cases and controls were compared using two sample t-tests for continuous variables and Chi-square or Fisher’s exact tests for categorical variables where appropriate.

The metabolic feature intensities were summarized as median values of the non-zero readings across triplicates. Features were retained for analysis if there were less than 20% zero readings across CD samples and control samples. Intensities were log 2 (intensity+1) transformed, mean centered and scaled by standard deviation.

Logistic regression model was utilized for the metabolome-wide association study (MWAS). The association was examined between the binary response variable of CD and each of the metabolic features, including age, gender, diet restrictions, and BMI as covariates. The Benjamini-Hochberg false discovery rate (FDR) [24] method was used to correct for multiple comparisons. Analyses were performed using R version 3.3.3 (R Foundation for Statistical Computing, Vienna, Austria) and SAS statistical software version 9.4 (SAS Institute Inc, Cary, NC).

### Pathway Analysis

Features associated with CD that had p<0.05 in MWAS were chosen as target features that entered pathway analysis in Mummichog (version 2.1.1) [25], which is a software designed for high throughput untargeted metabolomics data. This analysis allows us to explore the functional activities directly from the untargeted features, bypassing metabolite identification. All features entered MWAS were utilized as the feature reference pool. Pathways enriched with p<0.05 were considered as significant.

## Results

Among the 200 subjects, 100 were CD cases and 100 were controls (Table 1). The proportion of females was significantly higher among cases than controls (78% vs 55%, p<0.001). Cases were older compared to controls at enrollment (62.3 vs 57.5, p=0.011), and had lower BMI (26.1 vs 28.4, p=0.002). All of these variables were adjusted in the statistical model as covariates.

**Table 1.** Baseline characteristics.

| Variable | Level | Control N=100 | CD N=100 | P-value<sup>*</sup> |
| --- | --- | --- | --- | --- |
| Female (%) |  | 55 (55) | 78 (78) | **<.001** |
| Ethnicity | Black (%) | 10 (10) | 4 (4) | 0.179 |
|  | White (%) | 88 (88) | 95 (95) |  |
|  | Other (%) | 2 (2) | 1 (1) |  |
| Age in year (sd) |  | 57.5 (14.6) | 62.3 (11.9) | **0.011** |
| BMI in kg/m<sup>2</sup> (sd) |  | 28.4 (6.0) | 26.1 (4.4) | **0.002** |
| Weight in lb (sd) |  | 181.4 (48.4) | 162.2 (34.7) | **0.001** |
| Height in inch (sd) |  | 67.1 (4.0) | 66.2 (3.8) | 0.074 |
| Dietary restrictions (%) |  | 9 (9) | 15 (15) | 0.277 |
| Smoking | Current smoker (%) | 7 (7) | 7 (7) | 0.912 |
|  | Past smoker (%) | 27 (27) | 30 (30) |  |
|  | Never smoker (%) | 64 (64) | 62 (62) |  |

\*Two-sample t-tests used for continuous variables; Chi-square tests or Fisher's exact tests used for categorical variables where appropriate.

A total of 7,346 metabolic features passed quality control and entered MWAS. The MWAS revealed metabolomic associations for 289 features at p<0.05, and 35 features at p<0.01, and 1 feature at p<0.001. None of these features were significant at FDR cutoff 0.2 (Figure 1 & Supplemental Figure 1). Of the 289 features with p<0.05, 64 features were annotated based on Human Metabolome Database (Supplemental Table 1). Table 2 presents the 6 annotated features out of the 35 with p value < 0.01. The 289 features associated with CD at p<0.05 were further confirmed using the in-house library of confirmed metabolites. A total of 5 features were successfully matched (Table 3). Among those, higher level of uracil was associated with 36% lower odds of CD (m/z 111.0199, time 13.4s, OR 0.64, 95% CI 0.46-0.87, p=0.005).

**Table 2.**
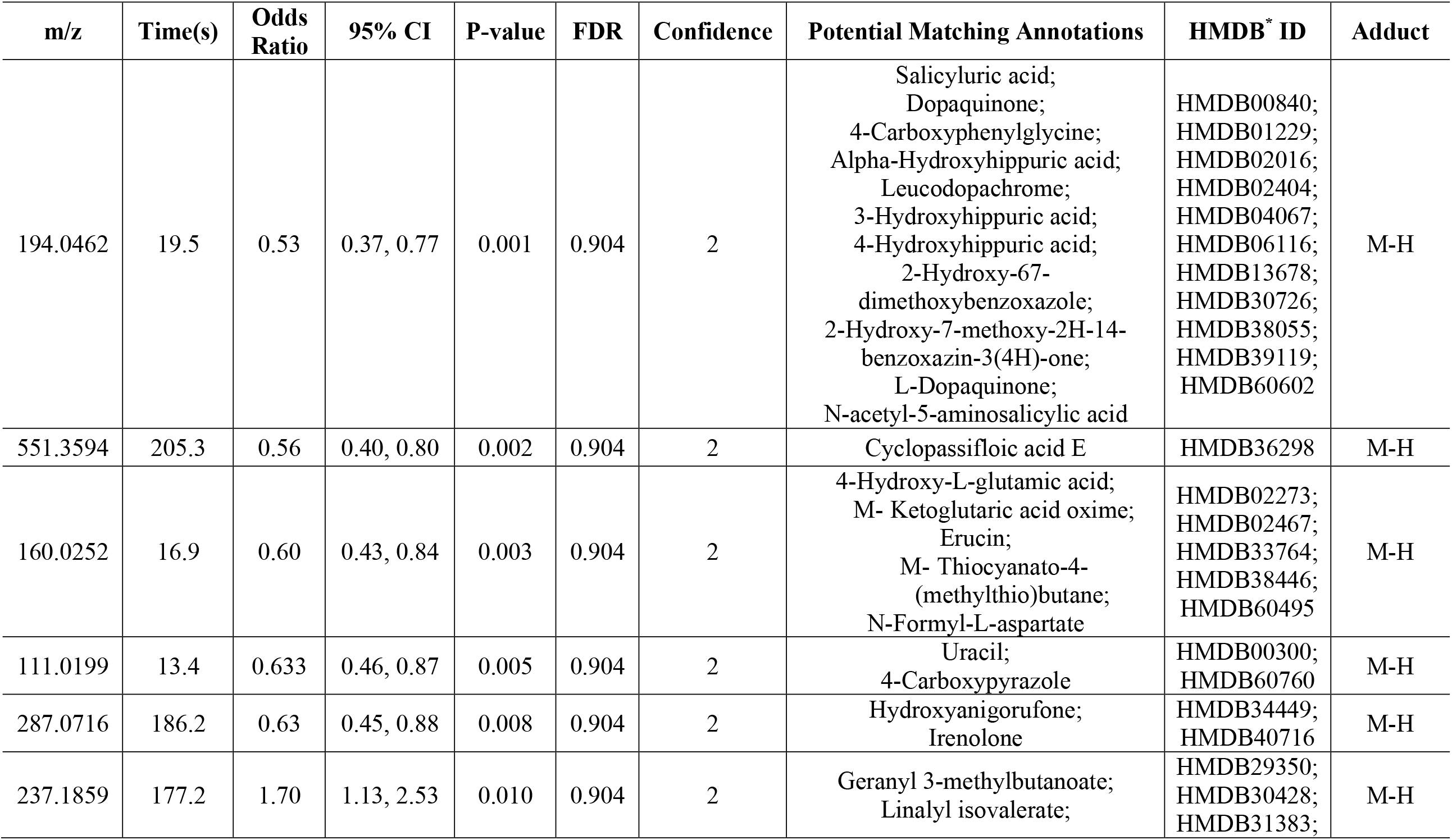
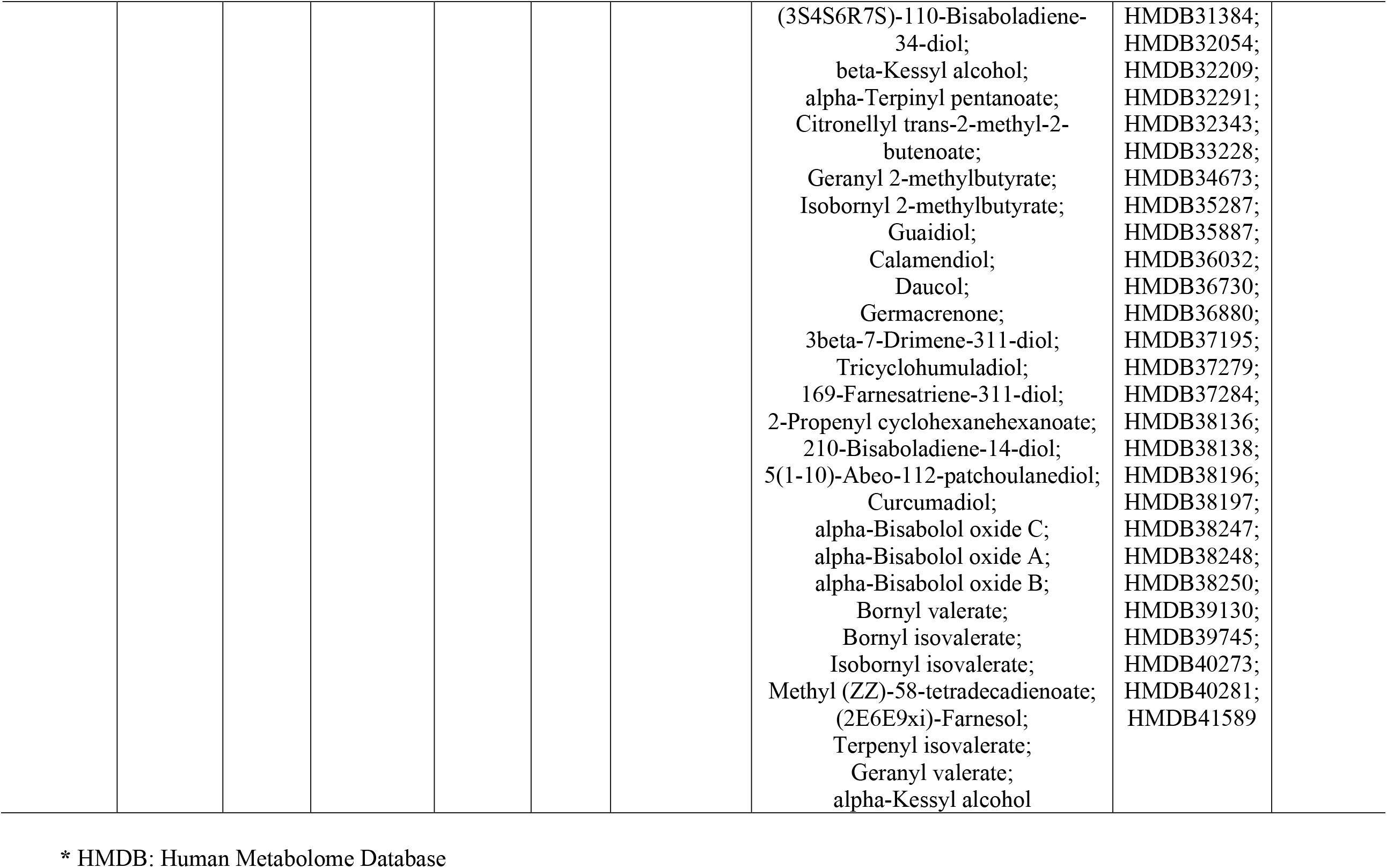
Metabolic features matched to possible annotations significantly associated with CD in MWAS at p<0.01.

**Table 3.**
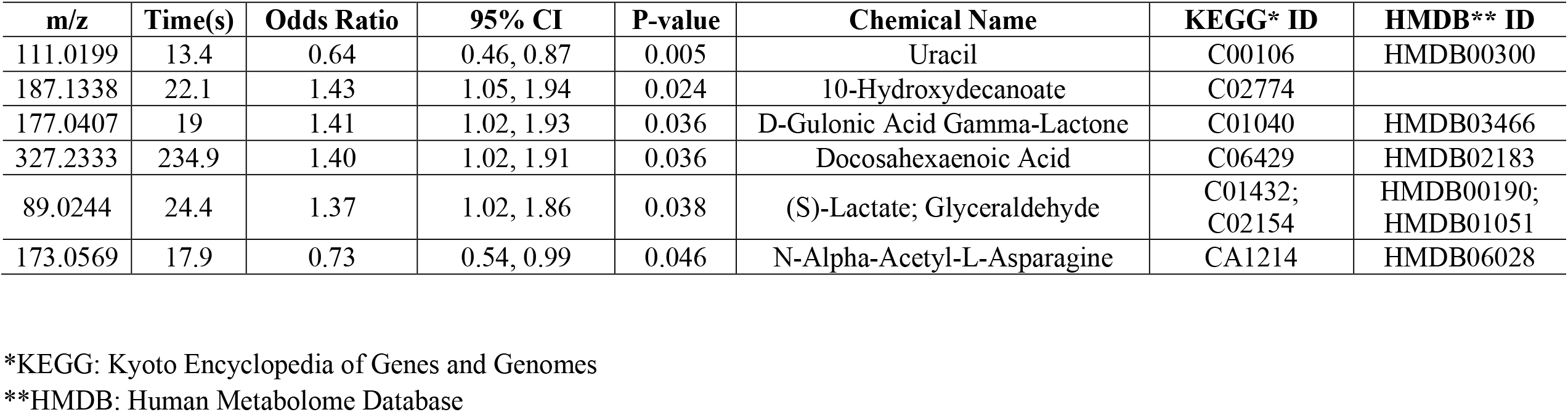
Confirmed features significantly associated with CD in MWAS at p<0.05.

**Figure 1.**
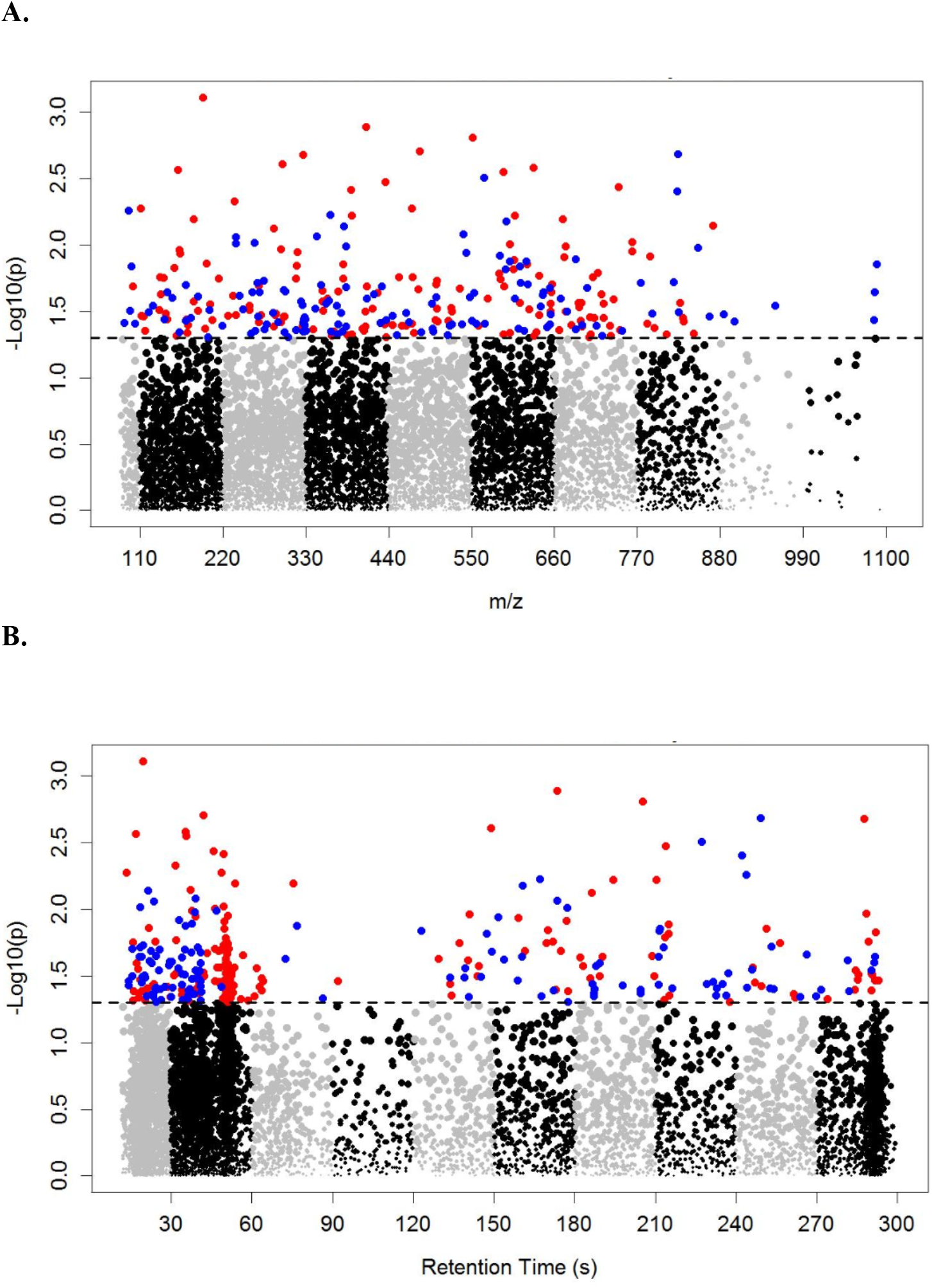
Manhattan plot of the metabolome-wide association study of CD. Total of 7,346 features. Dashed line indicates p=0.05. 157 features with p<0.05 and OR<1 colored in red; 132 features with p<0.05 and OR>1 colored in blue.

**Figure 2.**
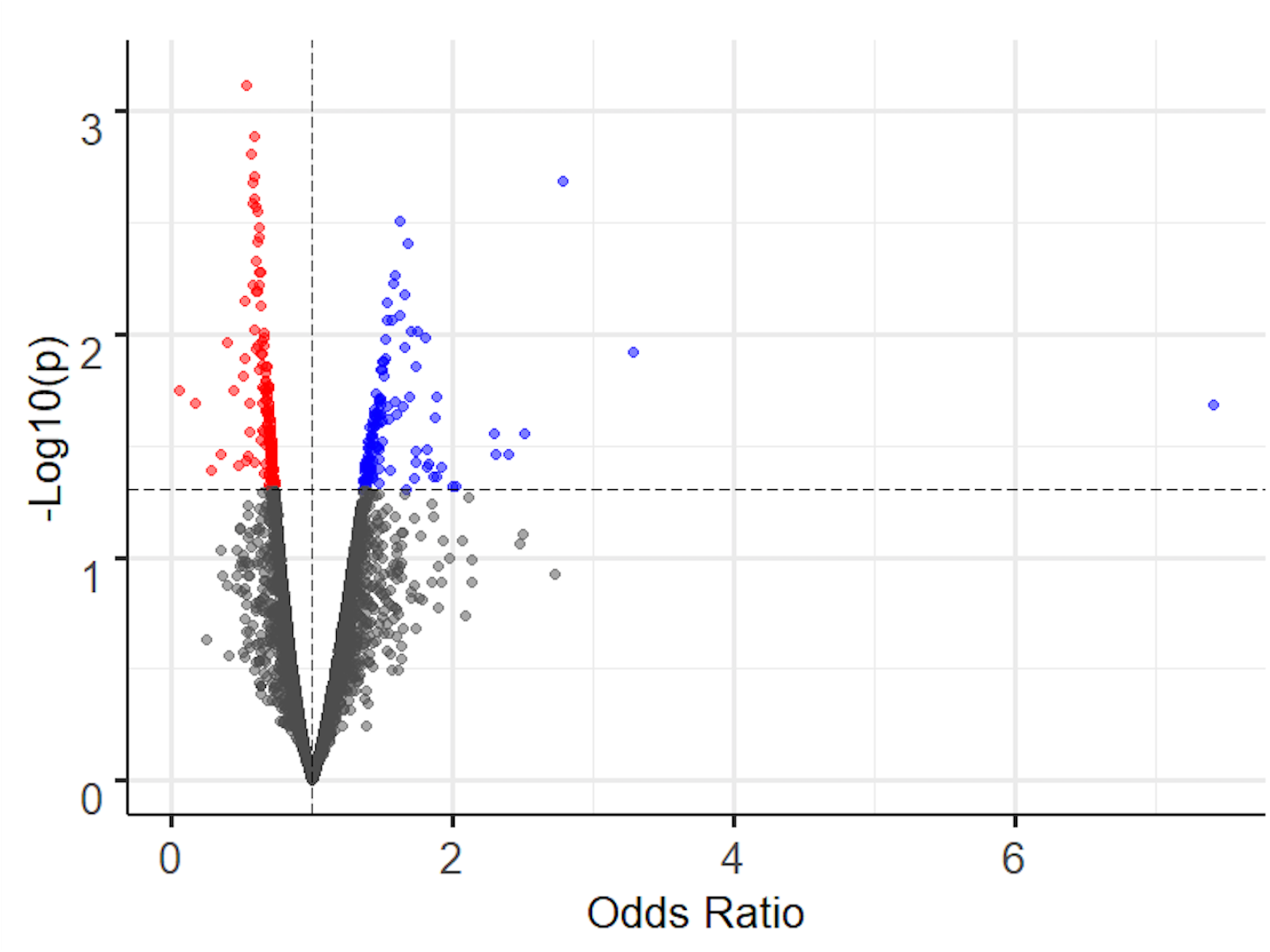
Volcano plot of the metabolome-wide association study of CD. Total of 7,346 features. Horizontal dashed line indicates p=0.05. 157 features with p<0.05 and OR<1 colored in red; 132 features with p<0.05 and OR>1 colored in blue.

Pathway analysis identified 9 significant pathways at p<0.05 that are enriched based on the 289 CD-associated metabolic features, including prostaglandin formation from dihomo gama-linoleic acid (p<0.001), pentose phosphate pathway (p=0.005), arachidonic acid metabolism (p=0.013), leukotriene metabolism (p=0.015), starch and sucrose metabolism (p=0.030), valine, leucine and isoleucine degradation (p=0.031), galactose metabolism (p=0.031), fructose and mannose metabolism (p=0.047) and propanoate metabolism (p=0.047) (Table 4).

**Table 4.** Significant pathways based on the 289 features associated with CD at p<0.05 in MWAS.

| **Pathways** | **Overlap size** | **Pathway size** | **P-value** |
| --- | --- | --- | --- |
| Prostaglandin formation from dihomo gamma-linoleic acid | 3 | 5 | <.001 |
| Pentose phosphate pathway | 4 | 22 | 0.005 |
| Arachidonic acid metabolism | 3 | 17 | 0.013 |
| Leukotriene metabolism | 3 | 18 | 0.015 |
| Starch and Sucrose Metabolism | 2 | 10 | 0.030 |
| Valine, leucine and isoleucine degradation | 3 | 23 | 0.031 |
| Galactose metabolism | 3 | 23 | 0.031 |
| Fructose and mannose metabolism | 2 | 14 | 0.047 |
| Propanoate metabolism | 2 | 14 | 0.047 |

## Discussion

This report describes the first large-scale metabolomics study for CD, the most common form of the idiopathic adult-onset dystonia. A total of 7,346 metabolites were measured, and significant differences could be detected between CD and control samples for individual metabolites. Pathway analysis revealed 9 biological processes to be associated at p<0.05. These results provide novel insights into the biological basis for CD.

An interesting metabolite that was identified as potentially abnormal in CD was the oxidized lipid 2-hydroxydecanoate. This metabolite was also identified in patients with Parkinson disease associated with mutations in *parkin* [26]. Dystonia is a well-known feature of this genetic subtype of Parkinson disease, although the dystonia is mostly in the limbs [27-29]. Docosahexaenoic acid was also potentially abnormal in CD. This metabolite is an omega-3 fatty acid which is essential for brain development and normal function, and many studies have implied that dystonia is a disorder of neural development or maladaptive plasticity [30, 31].

Among the biological pathways that were significantly associated with CD (Table 4), five were related to carbohydrate metabolism, suggesting a potential link with energy production. Three pathways were related to lipid metabolism, which produces prostaglandins and other metabolites known to mediate inflammatory responses [32, 33]. Involvement of these pathways may suggest a link to an immune-related process in dystonia. Several case series have reported CD to be associated with autoimmune diseases such as systemic lupus erythematosus, myasthenia gravis and Sjogrens syndrome [34-37]. CD can also occur as a delayed consequence of various infections, due to immune system cross-reactivity between the infectious agent and host targets [38-40]. Thus, the immune link suggested by our metabolomics findings may be worthy of further investigation.

Although this study has revealed several intriguing novel potential biological leads for future studies, the results must be interpreted with caution. Numerous variables are known to affect the metabolome in human blood samples, and it is not feasible to control for all of them. For instance, treatments for CD such as botulinum toxin injections may affect the metabolome. The timing of these injections is difficult to be incorporated in the statistical model due to missing data, and the dosing pattern varies individually. Although this study was based on a relatively large number of cases and controls all collected in a standardized manner at a single center, a replication study is still warranted. Because of the large number of measurements made, correction for multiple comparisons is needed to increase confidence in the findings. On the other hand, because this study was exploratory, the application of overly conservative statistical criteria is not helpful either, because important novel leads may be inappropriately overlooked. We therefore chose to report findings at different statistical thresholds. Additionally, due to the cross-sectional study design, no temporality between the different metabolomic profiles and CD can be inferred. A longitudinal cohort study with follow-up may be needed to better understand the disease onset and progression. Finally, using blood samples to infer biological mechanisms for a brain disorder carries significant limitations, given that relevant biomarkers may not cross the blood-brain barrier. Further investigations of cerebral spinal fluid or brain samples may be needed in future studies.

## Acknowledgements

This work was supported by the National Institute for Neurological Disorders and Stroke (NS096455 and NS116025) and the Rare Diseases Clinical Research Network of the Office of Rare Diseases Research at the National Center for Advancing Translational Sciences (TR001456).

## Author’s Roles

YVS and HAJ were involved in the conception, organization and execution of the research project;, LS, GK, AC, SF, AF and HAJ contributed to evaluation of cases, and phenotypic data collection and biospecimen collection; KU and DJ were responsible for the generation of metabolomic data; CL, KU and YVS were responsible of the design and execution of statistical analysis; The first draft was written by CL, HAJ and YVS, and developed further by LS, and GK. All co-authors contributed to critical review and editing of the manuscript.

