## Supplemental Table 1 for "A Metabolomic Study of Cervical Dystonia"

**Supplemental Table 1. Metabolic features matched to possible annotations significantly associated with CD in MWAS at $p<0.05$.**

| m/z | Time(s) | Odds Ratio | 95% CI | P-value | FDR | Confidence* | Potential Matching Annotations | HMDB** ID | Adduct |
|---|---|---|---|---|---|---|---|---|---|
| 194.0462 | 19.5 | 0.53 | 0.37, 0.77 | 0.001 | 0.904 | 2 | Salicyluric acid | HMDB00840 | M-H |
| | | | | | | | Dopaquinone | HMDB01229 | |
| | | | | | | | 4-Carboxyphenylglycine | HMDB02016 | |
| | | | | | | | Alpha-Hydroxyhippuric acid | HMDB02404 | |
| | | | | | | | Leucodopachrome | HMDB04067 | |
| | | | | | | | 3-Hydroxyhippuric acid | HMDB06116 | |
| | | | | | | | 4-Hydroxyhippuric acid | HMDB13678 | |
| | | | | | | | 2-Hydroxy-67-dimethoxybenzoxazole | HMDB30726 | |
| | | | | | | | 2-Hydroxy-7-methoxy-2H-14-benzoxazin-3(4H)-one | HMDB38055 | |
| | | | | | | | L-Dopaquinone | HMDB39119 | |
| | | | | | | | N-acetyl-5-aminosalicylic acid | HMDB60602 | |
| 551.3594 | 205.3 | 0.56 | 0.40, 0.80 | 0.002 | 0.904 | 2 | Cyclopassifloic acid E | HMDB36298 | M-H |
| 160.0252 | 16.9 | 0.60 | 0.43, 0.84 | 0.003 | 0.904 | 2 | 4-Hydroxy-L-glutamic acid | HMDB02273 | M-H |
| | | | | | | | A-Ketoglutaric acid oxime | HMDB02467 | |
| | | | | | | | Erucin | HMDB33764 | |
| | | | | | | | 1-Thiocyanato-4-(methylthio)butane | HMDB38446 | |
| | | | | | | | N-Formyl-L-aspartate | HMDB60495 | |
| 111.0199 | 13.4 | 0.64 | 0.46, 0.87 | 0.005 | 0.904 | 2 | Uracil | HMDB00300 | M-H |
| | | | | | | | 4-Carboxypyrazole | HMDB60760 | |
| 287.0716 | 186.2 | 0.63 | 0.45, 0.88 | 0.008 | 0.904 | 2 | Hydroxyanigorufone | HMDB34449 | M-H |
| | | | | | | | Irenolone | HMDB40716 | |
| 237.1859 | 177.2 | 1.70 | 1.14, 2.53 | 0.010 | 0.904 | 2 | Geranyl 3-methylbutanoate | HMDB29350 | M-H |
| | | | | | | | Linalyl isovalerate | HMDB30428 | |
| | | | | | | | (3S4S6R7S)-110-Bisaboladiene-34-diol | HMDB31383 | |
| | | | | | | | beta-Kessyl alcohol | HMDB31384 | |
| | | | | | | | alpha-Terpinyl pentanoate | HMDB32054 | |
| | | | | | | | Citronellyl trans-2-methyl-2-butenoate | HMDB32209 | |
| | | | | | | | Geranyl 2-methylbutyrate | HMDB32291 | |
| | | | | | | | Isobornyl 2-methylbutyrate | HMDB32343 | |
| | | | | | | | Guaidiol | HMDB33228 | |
| | | | | | | | Calamendiol | HMDB34673 | |
| | | | | | | | Daucol | HMDB35287 | |
| | | | | | | | Germacrenone | HMDB35887 | |
| | | | | | | | 3beta-7-Drimene-311-diol | HMDB36032 | |
| | | | | | | | Tricyclohumuladiol | HMDB36730 | |
| | | | | | | | 169-Farnesatriene-311-diol | HMDB36880 | |
| | | | | | | | 2-Propenyl cyclohexanehexanoate | HMDB37195 | |
| | | | | | | | 210-Bisaboladiene-14-diol | HMDB37279 | |

| | | | | | | | | | |
|---|---|---|---|---|---|---|---|---|---|
| | | | | | | | 5(1-10)-Abeo-112-patchoulanediol | HMDB37284 | |
| | | | | | | | Curcumadiol | HMDB38136 | |
| | | | | | | | alpha-Bisabolol oxide C | HMDB38138 | |
| | | | | | | | alpha-Bisabolol oxide A | HMDB38196 | |
| | | | | | | | alpha-Bisabolol oxide B | HMDB38197 | |
| | | | | | | | Bornyl valerate | HMDB38247 | |
| | | | | | | | Bornyl isovalerate | HMDB38248 | |
| | | | | | | | Isobornyl isovalerate | HMDB38250 | |
| | | | | | | | Methyl (ZZ)-58-tetradecadienoate | HMDB39130 | |
| | | | | | | | (2E6E9xi)-Farnesol | HMDB39745 | |
| | | | | | | | Terpenyl isovalerate | HMDB40273 | |
| | | | | | | | Geranyl valerate | HMDB40281 | |
| | | | | | | | alpha-Kessyl alcohol | HMDB41589 | |
| 163.0398 | 159.2 | 0.60 | 0.40, 0.89 | 0.012 | 0.904 | 2 | Phenylpyruvic acid | HMDB00205 | M-H |
| | | | | | | | m-Coumaric acid | HMDB01713 | |
| | | | | | | | 4-Hydroxycinnamic acid | HMDB02035 | |
| | | | | | | | 2-Hydroxycinnamic acid | HMDB02641 | |
| | | | | | | | Enol-phenylpyruvate | HMDB12225 | |
| | | | | | | | cis-p-Coumaric acid | HMDB30677 | |
| | | | | | | | 2-Oxo-3-phenylpropanoic acid (Mixture oxo and keto) | HMDB31629 | |
| | | | | | | | Coumaric acid | HMDB41592 | |
| 786.4774 | 177.1 | 0.63 | 0.44, 0.91 | 0.012 | 0.904 | 3 | Majonoside R2 | HMDB40418 | M-H_[-1] |
| | | | | | | | Vinaginsenoside R11 | HMDB40781 | |
| 318.0854 | 170.1 | 0.63 | 0.43, 0.91 | 0.014 | 0.904 | 2 | Ibandronate | HMDB14848 | M-H |
| 99.0813 | 122.9 | 1.49 | 1.08, 2.06 | 0.014 | 0.904 | 2 | 3-Hexanone | HMDB00753 | M-H |
| | | | | | | | 4-Methylpentanal | HMDB01318 | |
| | | | | | | | Methyl isobutyl ketone | HMDB02939 | |
| | | | | | | | 2-Oxohexane | HMDB05842 | |
| | | | | | | | Ethyl isopropyl ketone | HMDB05846 | |
| | | | | | | | Hexanal | HMDB05994 | |
| | | | | | | | (E)-3-Hexen-1-ol | HMDB30003 | |
| | | | | | | | 2-Hexen-1-ol | HMDB30952 | |
| | | | | | | | 2-Ethylbutanal | HMDB31220 | |
| | | | | | | | (E)-4-Hexen-1-ol | HMDB31502 | |
| | | | | | | | 1-Hexen-3-ol | HMDB31503 | |
| | | | | | | | 2-Methylpentanal | HMDB31578 | |
| 379.2863 | 256.3 | 0.44 | 0.23, 0.87 | 0.018 | 0.904 | 2 | MG(0:020:3(11Z14Z17Z)0:0) | HMDB11545 | M-H |
| | | | | | | | MG(0:020:3(5Z8Z11Z)0:0) | HMDB11546 | |
| | | | | | | | MG(0:020:3(8Z11Z14Z)0:0) | HMDB11547 | |
| | | | | | | | MG(20:3(11Z14Z17Z)0:00:0) | HMDB11575 | |
| | | | | | | | MG(20:3(5Z8Z11Z)0:00:0) | HMDB11576 | |
| | | | | | | | MG(20:3(8Z11Z14Z)0:00:0) | HMDB11577 | |
| | | | | | | | Isopersin | HMDB32735 | |

| | | | | | | | | | |
|---|---|---|---|---|---|---|---|---|---|
| | | | | | | | 2-Hydroxy-4-oxo-512-heneicosadien-1-yl acetate | HMDB39403 | |
| | | | | | | | Persin | HMDB41103 | |
| 317.0819 | 169.4 | 0.06 | 0.01, 0.61 | 0.018 | 0.904 | 2 | Musanolone F | HMDB41451 | M-H |
| 274.1055 | 19.6 | 1.45 | 1.06, 1.98 | 0.019 | 0.904 | 2 | Norophthalmic acid | HMDB05766 | M-H |
| | | | | | | | Gamma-Glutamyl Glutamine | HMDB11738 | |
| 818.5888 | 253 | 1.89 | 1.11, 3.22 | 0.019 | 0.904 | 3 | PS(18:020:0) | HMDB10164 | M-H |
| 624.1299 | 15.5 | 1.47 | 1.06, 2.04 | 0.020 | 0.904 | 3 | Quercetin 7-glucuronide 3-rhamnoside | HMDB36264 | M-H_[-1] |
| | | | | | | | Luteolin 4-glucoside 7-galacturonide | HMDB38809 | |
| 350.21 | 35.2 | 1.58 | 1.08, 2.33 | 0.020 | 0.904 | 2 | Sphingosine 1-phosphate (d16:1-P) | HMDB60061 | M-H |
| 101.0243 | 161.6 | 0.65 | 0.45, 094 | 0.021 | 0.904 | 2 | 2-Ketobutyric acid | HMDB00005 | M-H |
| | | | | | | | Acetoacetic acid | HMDB00060 | |
| | | | | | | | 2-Methyl-3-oxopropanoic acid | HMDB01172 | |
| | | | | | | | Succinic acid semialdehyde | HMDB01259 | |
| | | | | | | | (S)-Methylmalonic acid semialdehyde | HMDB02217 | |
| | | | | | | | 4-Hydroxycrotonic acid | HMDB03381 | |
| | | | | | | | Acetic anhydride | HMDB31646 | |
| 301.2389 | 266.3 | 1.44 | 1.05, 1.97 | 0.022 | 0.904 | 3 | MG(0:014:00:0) | HMDB11530 | M-H |
| | | | | | | | MG(14:00:00:0) | HMDB11561 | |
| 373.0454 | 208.8 | 0.67 | 0.48, 0.94 | 0.022 | 0.904 | 2 | 6-Hydroxymethyletoricoxib | HMDB13997 | M-H |
| | | | | | | | Etoricoxib 1-N-oxide | HMDB60926 | |
| 303.0191 | 24.4 | 1.46 | 1.05, 2.02 | 0.023 | 0.904 | 2 | 5-(345-Trihydroxyphenyl)-gamma-valerolactone-3-O-sulphate | HMDB59985 | M-H |
| | | | | | | | 5-(345-Trihydroxyphenyl)-gamma-valerolactone-4-O-sulphate | HMDB59987 | |
| | | | | | | | Sinapinic acid-O-sulphate | HMDB60020 | |
| 656.151 | 190.3 | 0.69 | 0.50, 0.95 | 0.023 | 0.904 | 3 | Teniposide | HMDB14587 | M-H_[-1] |
| | | | | | | | Patuletin 3-gentiobioside | HMDB37541 | |
| 146.0552 | 291.6 | 1.44 | 1.05, 1.97 | 0.023 | 0.904 | 3 | 2-Methylglutaric acid | HMDB00422 | M-H_[-1] |
| | | | | | | | Adipic acid | HMDB00448 | |
| | | | | | | | Methylglutaric acid | HMDB00752 | |
| | | | | | | | Monomethyl glutaric acid | HMDB00858 | |
| | | | | | | | 22-Dimethylsuccinic acid | HMDB02074 | |
| | | | | | | | Solerol | HMDB02173 | |
| | | | | | | | (S)-2-Aceto-2-hydroxybutanoic acid | HMDB06900 | |
| | | | | | | | Dimethyl succinate | HMDB33837 | |
| 233.0667 | 140.5 | 0.70 | 0.52, 0.95 | 0.024 | 0.904 | 2 | 1-Isopropyl citrate | HMDB32438 | M-H |
| | | | | | | | 2-Isopropyl citrate | HMDB38083 | |
| 242.1765 | 281.4 | 1.54 | 1.06, 2.25 | 0.024 | 0.904 | 2 | N-Undecanoylglycine | HMDB13286 | M-H |
| 187.1338 | 22.1 | 1.43 | 1.05, 1.94 | 0.024 | 0.904 | 2 | 3-Hydroxycapric acid | HMDB02203 | M-H |
| | | | | | | | (R)-3-Hydroxydecanoic acid | HMDB10725 | |
| | | | | | | | 9-Hydroxydecanoic acid | HMDB33201 | |
| | | | | | | | (1R2R4R8R)-p-Menthane-289-triol | HMDB33574 | |
| | | | | | | | 7-Methyl-3-methylene-167-octanetriol | HMDB33640 | |

| | | | | | | | | | |
|---|---|---|---|---|---|---|---|---|---|
| | | | | | | | cis-p-Menthane-178-triol | HMDB34783 | |
| | | | | | | | 26-Dimethyl-7-octene-236-triol | HMDB38186 | |
| | | | | | | | (1S2S4R8R)-p-Menthane-129-triol | HMDB39470 | |
| | | | | | | | 2-Hexyl-13-dioxan-5-ol | HMDB39668 | |
| | | | | | | | 2-Hexyl-13-dioxolane-4-methanol | HMDB39669 | |
| | | | | | | | (1R2R4S)-p-Menthane-128-triol | HMDB39893 | |
| | | | | | | | xi-5-Hydroxydecanoic acid | HMDB40329 | |
| | | | | | | | Ethyl ()-3-hydroxyoctanoate | HMDB41600 | |
| 323.2593 | 187.9 | 1.42 | 1.04, 1.95 | 0.027 | 0.904 | 3 | (Z)-15-Oxo-11-eicosenoic acid | HMDB29797 | M-H |
| | | | | | | | (13R14R)-8-Labdene-131415-triol | HMDB34957 | |
| | | | | | | | (13R14R)-7-Labdene-131415-triol | HMDB34958 | |
| 203.0677 | 17.6 | 0.70 | 0.51, 0.96 | 0.028 | 0.904 | 2 | L-beta-aspartyl-L-alanine | HMDB11162 | M-H |
| | | | | | | | 5-L-Glutamylglycine | HMDB11667 | |
| | | | | | | | Alanyl-Aspartate | HMDB28683 | |
| | | | | | | | Aspartyl-Alanine | HMDB28746 | |
| 325.2389 | 245.9 | 1.42 | 1.04, 1.94 | 0.029 | 0.904 | 3 | 1-Acetoxy-2-hydroxy-16-heptadecen-4-one | HMDB31006 | M-H |
| | | | | | | | Avocadyne 2-acetate | HMDB31047 | |
| | | | | | | | Avocadyne 1-acetate | HMDB31048 | |
| | | | | | | | Avocadyne 4-acetate | HMDB31049 | |
| 371.0065 | 25.4 | 1.43 | 1.04, 1.96 | 0.029 | 0.904 | 3 | Sodium 6-hydroxy-5-(phenylazo)-2-naphthalenesulfoniate | HMDB32902 | M+Na-2H |
| 952.775 | 39 | 1.42 | 1.04, 1.94 | 0.029 | 0.904 | 2 | PC(24:1(15Z)24:1(15Z)) | HMDB08816 | M-H |
| 255.0667 | 284.5 | 0.71 | 0.52, 0.96 | 0.029 | 0.904 | 2 | Dihydrodaidzein | HMDB05760 | M-H |
| | | | | | | | (E)-244-Trihydroxychalcone | HMDB29462 | |
| | | | | | | | (2S)-Liquiritigenin | HMDB29519 | |
| | | | | | | | (S)-Pinocembrin | HMDB30808 | |
| | | | | | | | 4-Methoxybenzophenone-2-carboxylic acid | HMDB32577 | |
| | | | | | | | 7E-Mycosinyl acetate | HMDB33904 | |
| | | | | | | | Emodinanthranol | HMDB36457 | |
| | | | | | | | Isoliquiritigenin | HMDB37316 | |
| | | | | | | | 2-Dehydro-O-desmethylangolensin | HMDB41647 | |
| 255.1605 | 285.5 | 0.67 | 0.47, 0.96 | 0.031 | 0.904 | 2 | 5-Nonyltetrahydro-2-oxo-3-furancarboxylic acid | HMDB30993 | M-H |
| | | | | | | | Monomenthyl succinate | HMDB36143 | |
| 97.0294 | 20.1 | 1.43 | 1.03, 1.98 | 0.031 | 0.904 | 2 | 2-Furanmethanol | HMDB13742 | M-H |
| | | | | | | | 5-Methyl-2(3H)-furanone | HMDB29609 | |
| | | | | | | | 5-Hydroxy-4-pentenoic acid d-lactone | HMDB40145 | |
| 409.2728 | 209.5 | 0.71 | 0.52, 0.97 | 0.032 | 0.904 | 2 | 58-Epoxy-58-dihydro-10-apo-by-carotene-310-diol | HMDB39020 | M-H |
| | | | | | | | 56-Epoxy-56-dihydro-10-apo-by-carotene-310-diol | HMDB39021 | |
| 655.1483 | 189.3 | 0.71 | 0.52, 0.97 | 0.032 | 0.904 | 3 | Teniposide | HMDB14587 | M-H |
| | | | | | | | Patuletin 3-gentiobioside | HMDB37541 | |
| 121.0657 | 145.1 | 1.41 | 1.03, 1.93 | 0.032 | 0.904 | 2 | 4-Ethylphenol | HMDB29306 | M-H |
| | | | | | | | 25-Dimethylphenol | HMDB30540 | |
| | | | | | | | 1-Methoxy-2-methylbenzene | HMDB32074 | |
| | | | | | | | 1-Methoxy-4-methylbenzene | HMDB32076 | |

| | | | | | | | | | |
|---|---|---|---|---|---|---|---|---|---|
| | | | | | | | 23-Dimethylphenol | HMDB32148 | |
| | | | | | | | 26-Dimethylphenol | HMDB32150 | |
| | | | | | | | 34-Dimethylphenol | HMDB32151 | |
| | | | | | | | 1-Phenylethanol | HMDB32619 | |
| | | | | | | | 2-Phenylethanol | HMDB33944 | |
| | | | | | | | 4-Methylbenzyl alcohol | HMDB41609 | |
| | | | | | | | 3-Ethylphenol | HMDB59873 | |
| 824.6935 | 19.2 | 1.46 | 1.03, 2.06 | 0.032 | 0.904 | 2 | Glucosylceramide (d18:125:0) | HMDB04979 | M-H |
| 289.0875 | 285.3 | 0.71 | 0.51, 0.97 | 0.034 | 0.904 | 2 | N-gamma-Glutamyl-S-allylcysteine | HMDB31874 | M-H |
| | | | | | | | 23-Dihydro-23-dihydroxy-9-phenyl-1H-phenalen-1-one | HMDB34688 | |
| | | | | | | | N-gamma-Glutamyl-S-(1-propenyl)cysteine | HMDB38553 | |
| | | | | | | | N-gamma-Glutamyl-S-trans-(1-propenyl)cysteine | HMDB40557 | |
| 227.0928 | 291.5 | 0.71 | 0.52, 0.97 | 0.034 | 0.904 | 2 | Propyl propane thiosulfonate | HMDB32496 | M-H |
| 301.1813 | 46.2 | 0.72 | 0.53, 0.98 | 0.035 | 0.904 | 2 | 2-Hydroxyestradiol-3-methyl ether | HMDB00380 | M-H |
| | | | | | | | 2-Methoxyestradiol | HMDB00405 | |
| | | | | | | | 19-Hydroxyandrost-4-ene-317-dione | HMDB03955 | |
| | | | | | | | 19-Oxotestosterone | HMDB03959 | |
| | | | | | | | 7a-Hydroxyandrost-4-ene-317-dione | HMDB06771 | |
| | | | | | | | 11b-Hydroxyandrost-4-ene-317-dione | HMDB06773 | |
| | | | | | | | 16a-Hydroxyandrost-4-ene-317-dione | HMDB06774 | |
| | | | | | | | 4-Methoxy-17beta-estradiol | HMDB12782 | |
| | | | | | | | (2Z8S9Z)-29-Heptadecadiene-8-hydroxy-46-diyne-1-yl acetate | HMDB31038 | |
| | | | | | | | 8-Dehydroshogaol | HMDB33125 | |
| | | | | | | | 2-(37-Dimethyl-26-octadienyl)-4-hydroxy-6-methoxyacetophenone | HMDB34028 | |
| | | | | | | | Falcarindiol 3-acetate | HMDB34167 | |
| | | | | | | | Yucalexin B11 | HMDB36696 | |
| | | | | | | | Ginsenoyne H | HMDB39568 | |
| | | | | | | | 2-Hexyl-5-2-(4-hydroxy-3-methoxyphenyl)ethylfuran | HMDB40927 | |
| | | | | | | | Acetylpanaxydol | HMDB41204 | |
| 115.0399 | 92.1 | 0.70 | 0.50, 0.97 | 0.035 | 0.904 | 2 | Alpha-ketoisovaleric acid | HMDB00019 | M-H |
| | | | | | | | Methylacetoacetic acid | HMDB00310 | |
| | | | | | | | Levulinic acid | HMDB00720 | |
| | | | | | | | 2-Oxovaleric acid | HMDB01865 | |
| | | | | | | | 2-Methylacetoacetic acid | HMDB03771 | |
| | | | | | | | Glutarate semialdehyde | HMDB12233 | |
| | | | | | | | Ethyl pyruvate | HMDB31643 | |
| | | | | | | | Acetoxyacetone | HMDB34466 | |
| 267.0728 | 14.2 | 2.30 | 1.06, 5.00 | 0.035 | 0.904 | 2 | Inosine | HMDB00195 | M-H |
| | | | | | | | 3-Deoxy-D-glycero-D-galacto-2-nonulosonic acid | HMDB00425 | |

| | | | | | | | | | |
|---|---|---|---|---|---|---|---|---|---|
| | | | | | | | Allopurinol riboside | HMDB00481 | |
| | | | | | | | Arabinosylhypoxanthine | HMDB03040 | |
| 328.2365 | 231.5 | 1.40 | 1.02, 1.91 | 0.035 | 0.904 | 3 | Docosahexaenoic acid | HMDB02183 | M-H_[-1] |
| | | | | | | | Neogrifolin | HMDB30053 | |
| | | | | | | | Grifolin | HMDB30446 | |
| | | | | | | | Retinol acetate | HMDB35185 | |
| | | | | | | | (ZZ)-2-Methyl-5-(81114-pentadecatrienyl)-13-benzenediol | HMDB38908 | |
| 177.0407 | 19 | 1.41 | 1.02, 1.93 | 0.036 | 0.904 | 2 | Gluconolactone | HMDB00150 | M-H |
| | | | | | | | 2-Keto-3-deoxy-D-gluconic acid | HMDB01353 | |
| | | | | | | | 3-Keto-b-D-galactose | HMDB01385 | |
| | | | | | | | Galactonolactone | HMDB02541 | |
| | | | | | | | L-Gulonolactone | HMDB03466 | |
| | | | | | | | Melizame | HMDB29684 | |
| | | | | | | | D-Arabino-hexos-2-ulose | HMDB29932 | |
| | | | | | | | Methylthiomethyl 2-methylbutanethiolate | HMDB31714 | |
| 327.2333 | 234.9 | 1.40 | 1.02, 1.91 | 0.036 | 0.904 | 3 | Docosahexaenoic acid | HMDB02183 | M-H |
| | | | | | | | Neogrifolin | HMDB30053 | |
| | | | | | | | Grifolin | HMDB30446 | |
| | | | | | | | Retinol acetate | HMDB35185 | |
| | | | | | | | (ZZ)-2-Methyl-5-(81114-pentadecatrienyl)-13-benzenediol | HMDB38908 | |
| 329.2369 | 229.2 | 1.39 | 1.02, 1.90 | 0.036 | 0.904 | 3 | Docosahexaenoic acid | HMDB02183 | M-H_[+2] |
| | | | | | | | Neogrifolin | HMDB30053 | |
| | | | | | | | Grifolin | HMDB30446 | |
| | | | | | | | Retinol acetate | HMDB35185 | |
| | | | | | | | (ZZ)-2-Methyl-5-(81114-pentadecatrienyl)-13-benzenediol | HMDB38908 | |
| 206.1037 | 23.2 | 0.53 | 0.29, 0.96 | 0.037 | 0.904 | 2 | Miglitol | HMDB14634 | M-H |
| 297.1348 | 28.6 | 0.67 | 0.45, 0.98 | 0.038 | 0.904 | 2 | Toxin T2 tetrol | HMDB36159 | M-H |
| | | | | | | | 37815-Scirpenetetrol | HMDB37560 | |
| 89.0244 | 24.4 | 1.37 | 1.02, 1.86 | 0.038 | 0.904 | 3 | L-Lactic acid | HMDB00190 | M-H |
| | | | | | | | Hydroxypropionic acid | HMDB00700 | |
| | | | | | | | Glyceraldehyde | HMDB01051 | |
| | | | | | | | D-Lactic acid | HMDB01311 | |
| | | | | | | | Dihydroxyacetone | HMDB01882 | |
| | | | | | | | Dimethyl carbonate | HMDB29580 | |
| | | | | | | | Monoethyl carbonate | HMDB31232 | |
| | | | | | | | Methoxyacetic acid | HMDB41929 | |
| 326.2423 | 244.1 | 1.39 | 1.02, 1.90 | 0.039 | 0.904 | 3 | 1-Acetoxy-2-hydroxy-16-heptadecen-4-one | HMDB31006 | M-H_[-1] |
| | | | | | | | Avocadyne 2-acetate | HMDB31047 | |
| | | | | | | | Avocadyne 1-acetate | HMDB31048 | |
| | | | | | | | Avocadyne 4-acetate | HMDB31049 | |
| 328.258 | 271.6 | 1.38 | 1.01, 1.87 | 0.040 | 0.904 | 3 | MG(0:016:1(9Z)0:0) | HMDB11534 | M-H_[-1] |
| | | | | | | | MG(16:1(9Z)0:00:0) | HMDB11565 | |
| | | | | | | | Avocadene 1-acetate | HMDB31043 | |

| | | | | | | | | | |
|---|---|---|---|---|---|---|---|---|---|
| | | | | | | | Avocadene 2-acetate | HMDB31044 | |
| | | | | | | | Avocadene 4-acetate | HMDB31045 | |
| 219.1026 | 172.1 | 1.40 | 1.01, 1.93 | 0.041 | 0.904 | 2 | 1-(5-Acetyl-2-hydroxyphenyl)-3-methyl-1-butanone | HMDB32589 | M-H |
| | | | | | | | Ethyl methyl-p-tolylglycidate | HMDB37492 | |
| | | | | | | | Ethyl 2-benzylacetoacetate | HMDB40424 | |
| 325.0541 | 177.5 | 0.73 | 0.54, 0.99 | 0.041 | 0.904 | 2 | Fertaric acid | HMDB29199 | M-H |
| 498.2628 | 187 | 1.40 | 1.01, 1.93 | 0.043 | 0.904 | 3 | LysoPE(0:020:5(5Z8Z11Z14Z17Z)) | HMDB11489 | M-H |
| | | | | | | | LysoPE(20:5(5Z8Z11Z14Z17Z)0:0) | HMDB11519 | |
| 263.1294 | 23.1 | 1.39 | 1.01, 1.92 | 0.044 | 0.904 | 2 | Hulupinic acid | HMDB30102 | M-H |
| | | | | | | | Heliespirone A | HMDB31972 | |
| | | | | | | | Curcolonol | HMDB33227 | |
| | | | | | | | 4-Hydroxydehydromyoporone | HMDB33661 | |
| | | | | | | | Isoamberboin | HMDB34719 | |
| | | | | | | | (S)-Abscisic acid | HMDB35140 | |
| | | | | | | | Tanacetin | HMDB35715 | |
| | | | | | | | Thellungianin G | HMDB35805 | |
| | | | | | | | (10R11R)-Pterosin L | HMDB35874 | |
| | | | | | | | 15-Hydroxymarasmen-3-one | HMDB36043 | |
| | | | | | | | O-Formyloreadone | HMDB36045 | |
| | | | | | | | Abscisic acid | HMDB36093 | |
| | | | | | | | (1beta8beta)-18-Dihydroxy-37(11)-eudesmadien-128-olide | HMDB36125 | |
| | | | | | | | Vulgarin | HMDB36130 | |
| | | | | | | | 3-Epiarmefolin | HMDB36135 | |
| | | | | | | | Alkhanin | HMDB36202 | |
| | | | | | | | Tavulin | HMDB36773 | |
| | | | | | | | Tatridin B | HMDB36931 | |
| | | | | | | | Istanbulin A | HMDB37055 | |
| | | | | | | | Blennin B | HMDB37528 | |
| | | | | | | | Umbellifolide | HMDB38156 | |
| | | | | | | | Enokipodin C | HMDB40119 | |
| | | | | | | | (8betaOH10beta)-8-Hydroxy-3-oxo-7(11)-eremophilen-128-olide | HMDB41227 | |
| | | | | | | | 2-Methyl-1-246-trihydroxy-3-(3-methyl-2-butenyl)phenyl-1-propanone | HMDB41284 | |
| 407.2572 | 215.3 | 0.73 | 0.54, 0.99 | 0.044 | 0.904 | 3 | Apo-10-violaxanthal | HMDB39018 | M-H |
| 330.2424 | 232.5 | 1.73 | 1.01, 2.94 | 0.044 | 0.904 | 3 | Docosahexaenoic acid | HMDB02183 | M-H_[+3] |
| | | | | | | | Neogrifolin | HMDB30053 | |
| | | | | | | | Grifolin | HMDB30446 | |
| | | | | | | | Retinol acetate | HMDB35185 | |
| | | | | | | | (ZZ)-2-Methyl-5-(81114-pentadecatrienyl)-13-benzenediol | HMDB38908 | |
| 327.2546 | 269.7 | 1.37 | 1.01, 1.85 | 0.045 | 0.904 | 3 | MG(0:016:1(9Z)0:0) | HMDB11534 | M-H |

| | | | | | | | | | |
|---|---|---|---|---|---|---|---|---|---|
| | | | | | | | MG(16:1(9Z)0:00:0) | HMDB11565 | |
| | | | | | | | Avocadene 1-acetate | HMDB31043 | |
| | | | | | | | Avocadene 2-acetate | HMDB31044 | |
| | | | | | | | Avocadene 4-acetate | HMDB31045 | |
| 499.2663 | 187 | 1.40 | 1.01, 1.94 | 0.045 | 0.904 | 3 | LysoPE(0:020:5(5Z8Z11Z14Z17Z)) | HMDB11489 | M-H_[-1] |
| | | | | | | | LysoPE(20:5(5Z8Z11Z14Z17Z)0:0) | HMDB11519 | |
| 302.2423 | 263.8 | 1.37 | 1.01, 1.86 | 0.045 | 0.904 | 3 | MG(0:014:00:0) | HMDB11530 | M-H_[-1] |
| | | | | | | | MG(14:00:00:0) | HMDB11561 | |
| 173.0569 | 17.9 | 0.73 | 0.54, 0.99 | 0.046 | 0.904 | 2 | Formiminoglutamic acid | HMDB00854 | M-H |
| | | | | | | | N-Acetylasparagine | HMDB06028 | |
| 473.3641 | 262 | 0.71 | 0.51, 0.99 | 0.046 | 0.904 | 3 | Soyasapogenol A | HMDB34505 | M-H |
| | | | | | | | Camelliagenin A | HMDB34528 | |
| | | | | | | | Priverogenin B | HMDB34644 | |
| | | | | | | | (3alphaOH20S24S)-319:2024-Diepoxydammarane-325-diol | HMDB34683 | |
| | | | | | | | Ganoderiol A | HMDB35326 | |
| | | | | | | | 2024-Epoxy-2526-dihydroxydammaran-3-one | HMDB39692 | |
| 385.26 | 213.4 | 0.74 | 0.54, 1.00 | 0.047 | 0.904 | 2 | Glycerol trihexanoate | HMDB31125 | M-H |
| | | | | | | | Mangalkanyl glucoside | HMDB36015 | |
| | | | | | | | Cryptomeridiol 11-rhamnoside | HMDB38018 | |
| 158.1017 | 15.9 | 0.73 | 0.53, 1.00 | 0.048 | 0.904 | 2 | ()-2-Pentylthiazolidine | HMDB40057 | M-H |

*Confidence: confidence score of the annotation, based on R package xMSannotator.
High confidence: score = 3. Medium confidence: score = 2. Low confidence: score < 2.

**HMDB: Human Metabolome Data Base.
